# Description of *Halopseudomonas maritima* sp. nov., based on phylogenomic analysis

**DOI:** 10.1101/2022.04.04.487078

**Authors:** Ritu Rani Archana Kujur, Subrata K Das

## Abstract

A novel bacterium, strain RR6^T^ of the genus *Halopseudomonas*, was isolated from the sea sand from Paradip port area, India. Cells are Gram-stain-negative, rod-shaped, aerobic, non-sporulating and motile. The major cellular fatty acids of RR6^T^ were C_10:0_ 3OH, C_12:0_, C_16:1_ w7c/16:1 w6c, 18:1 w7c and/or 18:1 w6c and C_16:0._ The predominant polar lipids were phosphatidylglycerol (PG), diphosphatidylglycerol (DPG), phosphatidylethanolamine (PE), phosphatidylcholine (PC), unidentified phospholipid (PL) and unidentified lipids (L1-L2). The genome size of the strain RR6 was 3.93 Mb. The genomic G+C content was 61.3%. Gene ontology study showed a significant fraction of genes were associated with biological processes (53%), followed by molecular function (36%) and cellular components (11%). Comparisons of 16S rRNA gene sequences revealed 99.73 - 99.87 % sequence similarity with the closely related type strains of the genus *Halopseudomonas*. The average nucleotide identity (ANI) and average amino acid identity (AAI) of RR6^T^ with reference strains of closely related species of the genus *Halopseudomonas* were below 95–96, and the corresponding *in-silico* DNA–DNA hybridization (DDH) values were below 70 %. A phylogenomic tree based on core genome analysis supported these results. Genotypic and phenotypic characteristics of RR6^T^ indicate that the strain represents a novel species of the genus *Halopseudomonas* and the name *Halopseudomonas maritima* sp. nov. is proposed. The type strain is RR6^T^ (= NBRC 115418^T^ = TBRC 15628^T^).

## INTRODUCTION

The family *Pseudomonadaceae* originally comprised the eighteen genera. Studies based on the evolutionary relationships among family *Pseudomonadaceae* introduced the genus *Halopseudomonas* by Rudra and Gupta [1]. At the time of writing, the genus *Halopseudomonas* comprises 14 species. *Halopseudomonas* have been isolated from various marine sources and desert sand, food waste, soil, air sample, aquatic plants, and algae intertidal shore [2-5]. Representatives of the genus *Halopseudomonas* Gram-negative, aerobic or facultatively anaerobic, non-spore-forming rods. Most species of this genus are motile by a single polar flagellum. The growth temperature ranges from 5 to 50 °C with optimal growth at 25–37 °C. The optimal pH for growth is between 6 and 10, and optimal growth occurs between 1 and 10 % (w/v) NaCl. The DNA G+C content is in the range of 57.5–63 mol%, and genome sizes range from 3.0 to 4.6 Mbp [1]. The present study describes a bacterium strain RR6^T^ isolated from sea sand collected from Paradip Port, Bay of Bengal. Its characterisation based on a polyphasic approach suggests it to be a new species of the genus *Halopseudomonas* for which the name *Halopseudomonas maritima* sp. nov. is proposed. The type strain is RR6^T^ (= TBRC 15628^T^ = NBRC 115418^T^), isolated from sea sand.

## ISOLATION AND ECOLOGY

Sand samples were collected in a sterile bottle from the seashore of Paradip port area (coordinates: 20°.16’36.3"N, 86°.40’02.1"E), India. Samples were transported to the laboratory, and the enrichment of the sand samples was carried out in marine agar broth 2216 (MA; Difco). After overnight incubation, aliquots (100µl) of the samples were serially diluted using PBS and plated onto marine agar 2216 (MA; Difco), and plates were incubated at 28 °C for three days. Few colonies that appeared on MA plates were picked and purified by repeated streaking on the same medium. A small, round, cream-colored colony designated strain RR6 was selected for further analysis. Cultures were maintained on marine agar (BD, Difco) and stored at 4 °C for short-term preservation. The cultures were stored at −80 °C in 15 % (v/v) glycerol for long-term preservation.

## GENOME SEQUENCE DETERMINATION

Genomic DNA was isolated by Xcelgen Bacterial gDNA Kit. The DNA sample was sheared to an average length of 15 kb using the Covaris system, as per the manufacturer’s protocol (Covaris, Woburn, MA, USA). Fragmented DNA was used for SMRTbell library preparation according to the manufacturer instruction. Quantity and quality of the SMRTbell libraries were evaluated using the High Sensitivity dsDNA kit and Qubit fluorometer and DNA 12,000 kit on the 2100 Bioanalyzer (Agilent, Santa Clara, CA, USA). The Whole-genome of *Halopseudomonas* strain RR6 was sequenced using the PacBio platform. A total of 610,251 high-quality reads were assembled using a canu-1.3 assembler (https://canu.readthedocs.io/en/latest/). The putative coding sequences (genes) were identified using the Prodigal-2.6.3 gene prediction tool (https://github.com/hyattpd/prodigal/releases/). The tRNA genes were identified using the tRNAscan-SE (v.1.31) tool, and rRNAs were identified using the RNAmmer 1.2 server [6-7]. The gene ontology (G.O.) study was performed using the Blastp program (https://www.biobam.com/omicsbox) implemented in Blast2GO tool [8] for cataloguing gene function. The genome size of the strain RR6 was 3.93 Mb. The genomic G+C content was 61.3%. The *de novo* assembly produced single scaffolds covering 3,935,609 bp (90.7%) with an average gene length of 1004 bp. The genome had 3, 616 genes, including 3532 protein-coding genes (CDS). The genome contains 55tRNA, three 5S rRNA, three 16S rRNA and three 23S rRNA. Gene ontology study showed a significant fraction of genes were associated with biological processes (53%), followed by molecular function (36%) and cellular components (11%) **(Fig. 1)**.

**Fig. 1.**
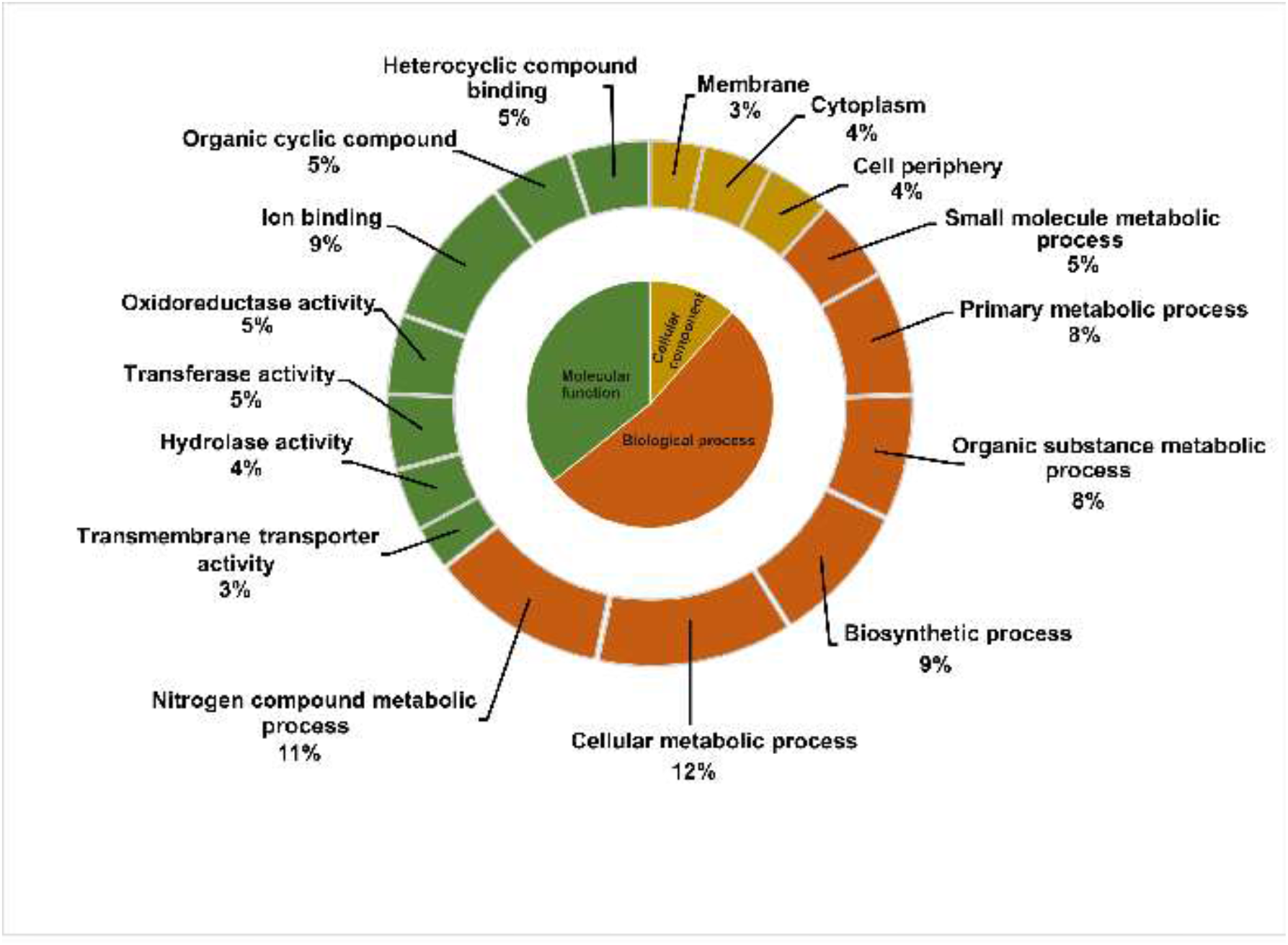
Gene ontology (GO) association of predicted protein-coding genes of *Halopseudomonas maritima* strain RR6.

## OVERALL GENOME RELATEDNESS INDICES

The average nucleotide identity (ANI) was calculated using the online tool [9]. *In silico* DDH similarity was measured with the help of the Genome-to-Genome Distance calculator (formula 2) [10]. Average amino acid identity (AAI) was estimated using the method of Rodriguez-R and Konstantinidis [11]. For the phylogenomic analysis using core genomes, presently available 13 whole-genome sequences of different *Halopseudomonas* species with more than 95% genome completeness were retrieved from the NCBI database using the NCBI-genome-download tool (https://github.com/kblin/ncbi-genome-download/). The core genes extracted by the UBCG pipeline [12] was concatenated, and a maximum-likelihood tree was reconstructed with the GTR model using the RAxML tool [13]. We compared the genomic relatedness of strain RR6 with reference genomes of *Halopseudomonas* available in the NCBI database (last accessed January 02, 2022). Now a days, whole-genome sequences information is essential for comparative phylogenetic and taxonomic studies of microorganisms. Different bioinformatics tools make it possible to compare genomic data of single organisms to determine the *in silico* DDH, ANI and AAI values. Thus, we evaluated the overall genome relatedness index (OGRI) to delineate the taxonomic and phylogenetic position of strain RR6. The ANI and AAI values between RR6 and the closely related reference species (https://lpsn.dsmz.de/genus/halopseudomonas; January 02, 2022) were below the threshold values (95-96%), justifying bacterial species delineation [14-15]. Thus, strain RR6 represents a novel species of the genus *Halopseudomonas*. Further, isDDH similarity values were less than 70 % to define bacterial species [10]. The ANI, AAI and isDDH data indicate strain RR6 could be proposed as a novel species of *Halopseudomonas* **(Table 1)**. Further, we analyzed the phylogenetic relationship between strain RR6 and reference strains based on 16S rRNA gene sequences and the genome-wide core genes. The maximum-likelihood (ML) phylogenetic tree showed strain RR6 clustered with *<u>Halopseudomonas gallaeciensis</u>* V113^T^ and *<u>Halopseudomonas pachastrellae</u>* CCUG 46540^T^ ; and formed a distinct clade (**Fig. 2**) which is similar to the phylogenetic tree based on the core genes **(Fig. 3)**, indicating the robustness of the tree topology.

**Table 1.** Comparison of the genomic characteristics of *Halopseudomonas* strain RR6 with closely related species of *Halopseudomonas*

| Sl. No. | Strain | Accession no. | Size (Mb) | G+C (%) | 16S rRNA Similarity (%) | *In silico* DDH (%) | ANIu (%) | AAI (%) |
| --- | --- | --- | --- | --- | --- | --- | --- | --- |
| 1 | *Halopseudomonas* strain RR6 | CP079801 | 3.93 | 61.3 | 100 | 100 | 100 | 100 |
| 2 | *Halopseudomonas gallaeciensis* V113<sup>T</sup> | LMAZ00000000 | 4.24 | 61.2 | 99.87 | 30.70 | 85.93 | 90.93 |
| 3 | *Halopseudomonas pachastrellae* CCUG 46540<sup>T</sup> | MUBC00000000 | 3.92 | 61.2 | 99.73 | 32.80 | 87.21 | 91.69 |

**Fig. 2.**
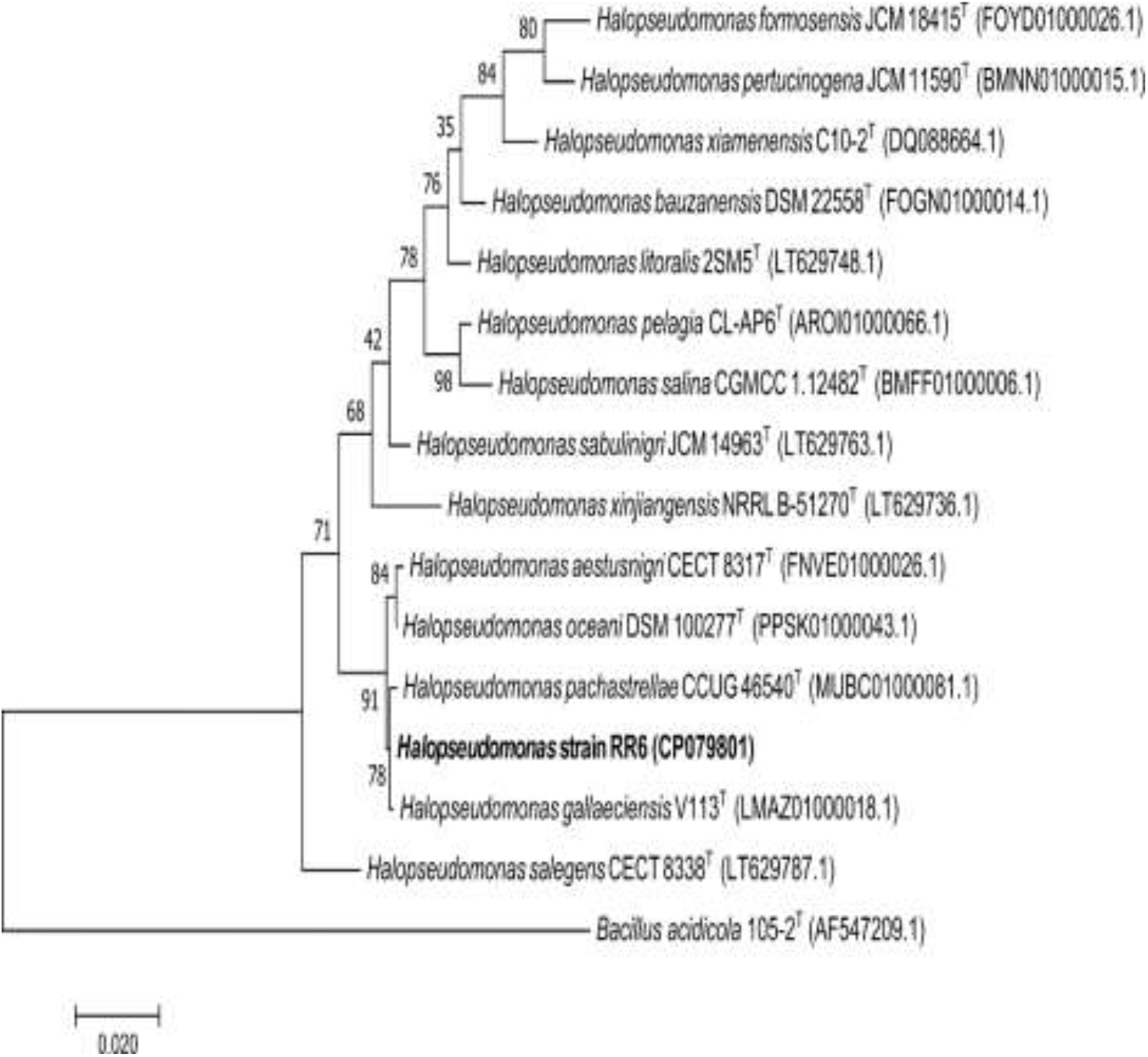
16S rRNA based maximum likelihood phylogenetic tree showing the position of *Halopseudomonas* strain RR6 among the related taxa. Bootstrap values expressed as percentages of 1000 replications are given at branch points. Accession numbers are given parentheses. Bar 2 substitution per 100 nucleotide position.

**Fig. 3.**
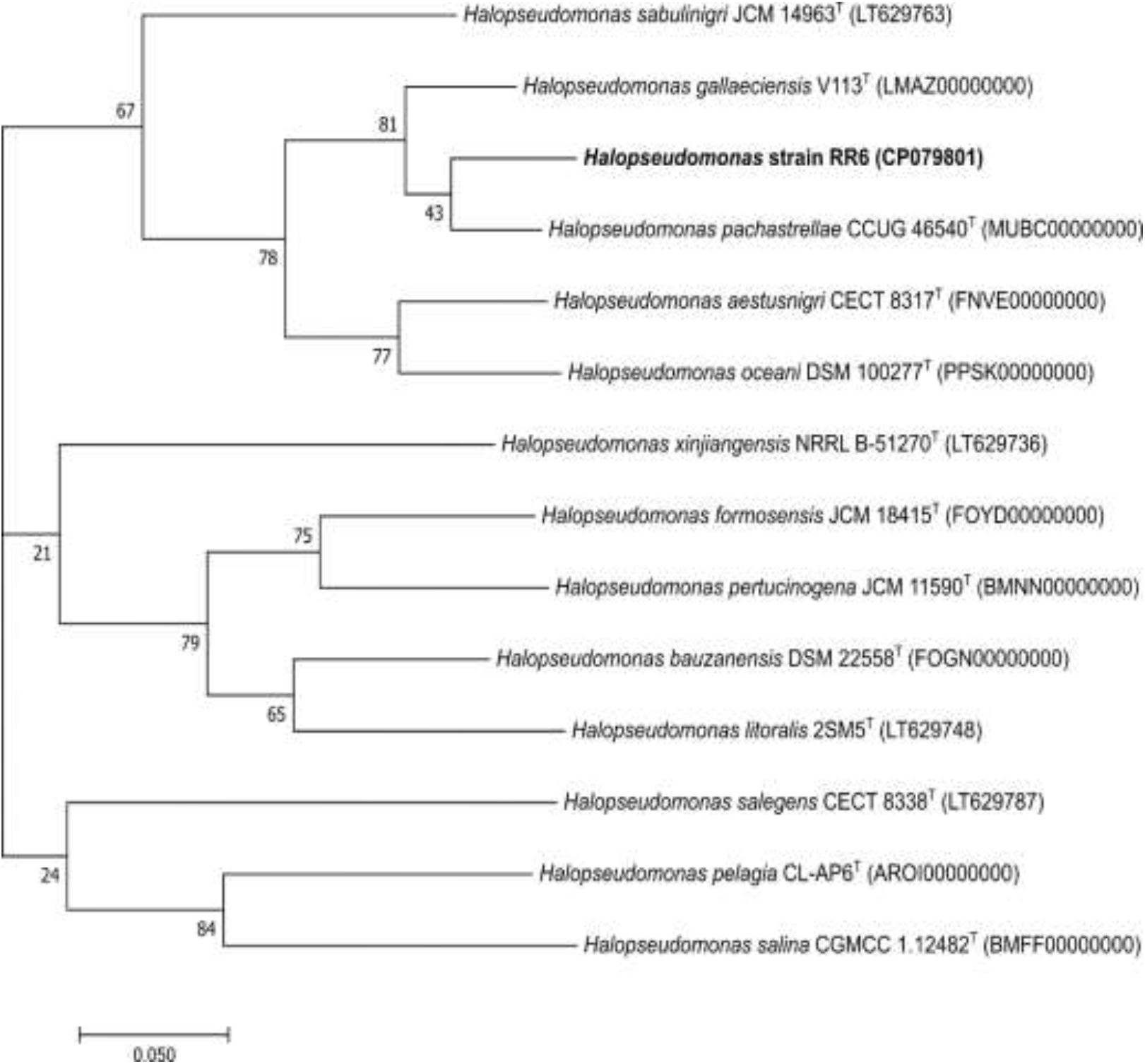
Phylogenetic tree based on the alignment of core genes of *Halopseudomonas* strain RR6 with 13 type strains (January 02, 2022).

## CHEMOTAXONOMIC CHARACTERIZATION

For the analysis of lipids, cells were grown in tryptic soy broth (Difco) to mid-exponential phase on a rotary shaker at 28 °C. Polar lipids were extracted from 200 mg of dry cells described by Bligh and Dyer [16] and separated by two-dimensional TLC (silica gel 60 F254, catalogue number 1.05554.0007; Merck). The solvent system used has been described earlier [17]. Cellular fatty acids were analyzed from the cells grown on trypticase soy agar (Difco) plates at 28°C for three days. Cells were saponified, and fatty acid methyl esters mixtures were separated using the Sherlock Microbial identification system (MIDI, Microbial ID), consisting of a model 6890 N gas chromatograph (Agilent). Polar lipids included phosphatidylglycerol (PG), diphosphatidylglycerol (DPG), phosphatidylethanolamine (PE), phosphatidylcholine (PC), unidentified phospholipid (PL) and unidentified lipids (L1-L2) (**Figure S1**). The lipid profile of strain RR6 was closely related to the reference strains of the genus *Halopseudomonas*. [5, 18]. Strain RR6 contained 10:0- 3OH (4.55%), 12:0 (8.65%), 16:1 w7c/16:1 w6c (15.86%), 18:1 w7c and/or 18:1 w6c (28.32%) and 16:0 (18.26%), as the dominant fatty acids (**Table S1**), in agreement with data previously reported for *<u>Halopseudomonas</u>*<u>type species</u>, justifying its placement in the genus *Halopseudomonas* [5, 19]. The overall chemotaxonomic characteristics of strain RR6 support its placement to the genus *Halopseudomonas*.

## PHENOTYPIC CHARACTERIZATION

The scanning electron microscope was used to study cell morphology (Carl Zeiss, EVO 18, Germany). Motility was tested using the hanging drop method [20]. Gram staining was done using a Gram staining kit (Becton-Dickinson, USA). Oxidase activity was assayed using commercial kit (Hi-Media, India). Catalase activity was assayed by mixing a culture pellet with a drop of hydrogen peroxide (10%, v/v). Anaerobic growth was tested using the BD GasPak EZ system (Becton Dickinson). Growth at different pH ranges (4.0–13.0; in steps of 0.5 pH units) and temperatures (4–50 °C; in steps of 2 °C) was determined in Marine broth. Growth at different temperatures was monitored by measuring OD_600_ with a spectrophotometer (CARY 300, UV–Vis spectrophotometer, Varian). The growth in different concentrations of NaCl (0–15 % (w/v) was determined in sea water medium (SWM) [18] and incubated at 28 °C for 2 days. Citrate utilization, activities of ornithine decarboxylase, urease, phenylalanine deaminase, the Voges–Proskauer test, hydrolysis of DNA, casein, starch, gelatin, arginine dihydrolase test, lysine decarboxylase test, production of H_2_S, indole production, the methyl red test, malonate utilization and nitrate reduction were assessed following the procedures described by Smibert & Krieg [21]. Phenotypic characterization was also performed using API 20 NE and API ZYM (bioMe’rieux). Cells are Gram-stain-negative, aerobic, non-sporulating, motile rod-shaped and 0.34–0.44 × 1.3 -1.6 μm in diameter (**Figure S2**). Different growth responses of the strain RR6 are shown in (**Table 2**).

**Table 2.**
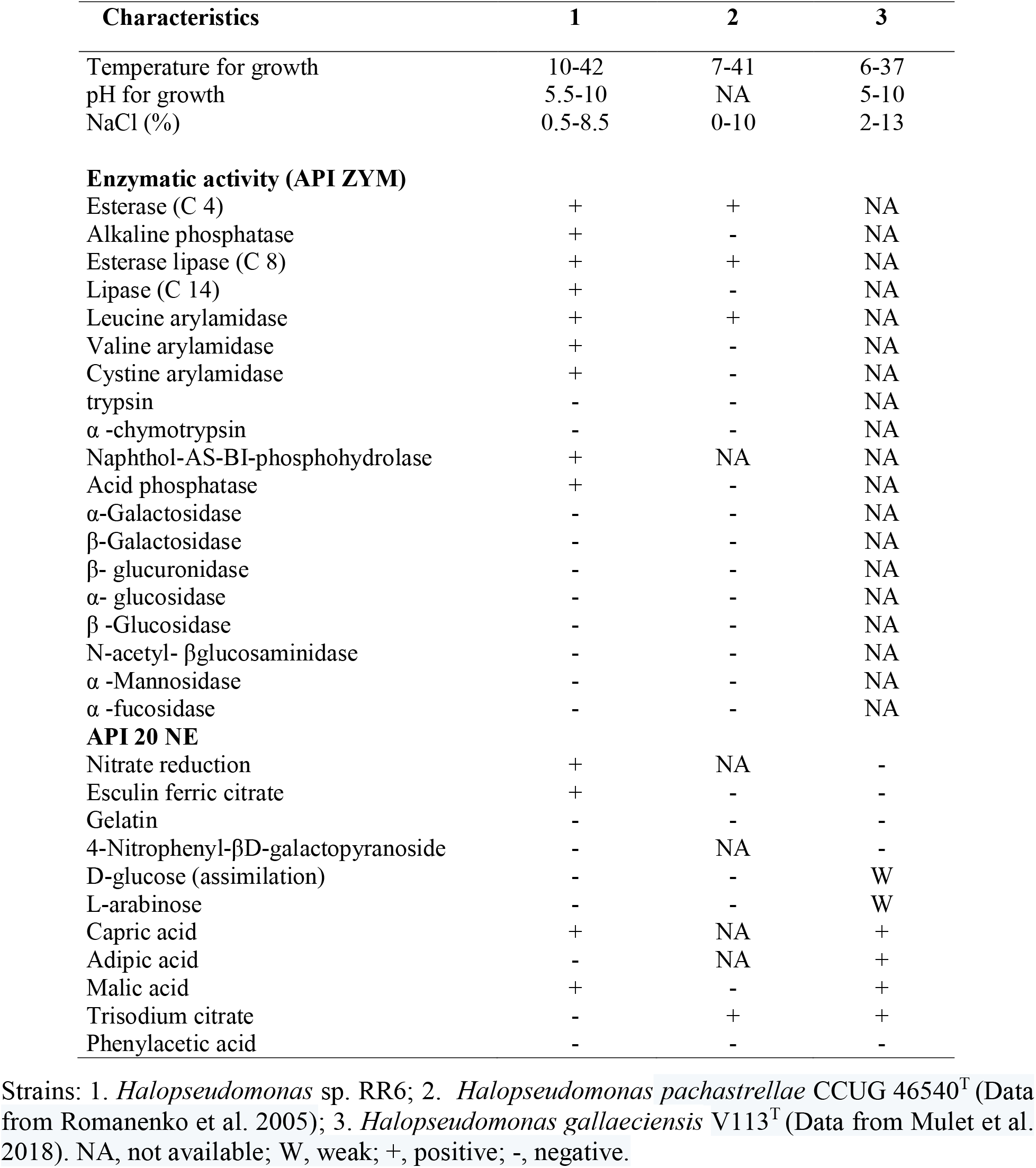
Metabolic and physiological characteristics of strain RR6 and closely related type species of the genus *Halopseudomonas*.

## DESCRIPTION OF *HALOPSEUDOMONAS MARITIMA* SP. NOV

*Halopseudomonas maritima* sp. nov. (ma.ri’ti.ma. L. fem. adj. *maritima*, belonging to the sea).

Cells are Gram-stain-negative, aerobic, non-sporulating, motile rod-shaped and 0.34–0.44 × 1.3 - 1.6 μm in diameter. Growth occurred at 10–42 °C and pH 5.5–10. Optimal growth was observed at 28–37°C and pH 6.0-8.0. It can grow at 0.5 -8.5 % (w/v) NaCl. Positive for lysine decarboxylase, catalase, malonate utilization, nitrate reduction and oxidase. Negative for citrate utilization, DNase, hydrolysis of casein, hydrolysis of starch, hydrolysis of gelatin, ornithine decarboxylase, arginine dihydrolase, Voges-Proskauer, methyl red, indole production, phenylalanine deaminase, urease and H_2_S production. In the API ZYM test, positive for esterase (C4), alkaline phosphatase, esterase lipase (C8), lipase (C14), leucine arylamidase, naphthol-AS-BI-phosphohydrolase, acid phosphatase, valine arylamidase and cystine arylamidase. In the API20NE test, positive for nitrate reduction, esculine, ferric citrate, capric acid and malic acid utilization. Predominant fatty acids are C_10:0_ 3OH, C_12:0_, C_16:1_ w7c/16:1 w6c, 18:1 w7c and/or 18:1 w6c and C_16:0._ Polar lipids include phosphatidylglycerol (PG), diphosphatidylglycerol (DPG), phosphatidylethanolamine (PE), phosphatidylcholine (PC), unidentified phospholipid (PL) and unidentified lipids (L1-L2).

The type strain is RR6^T^ (= TBRC 15628^T^ = NBRC 115418^T^), isolated from sea sand. The GenBank/EMBL/DDBJ accession numbers for the whole genome and 16S rRNA sequences of *Halopseudomonas maritima* strain RR6 are <u>MZ855216</u> and <u>CP079801</u>, respectively.

## Abbreviations

ANI,: average nucleotide identity;
AAI,: average amino acid identity;
isDDH,: *in silico* DNA-DNA hybridization.

## Acknowledgements

The author R.R.A.K. acknowledge the Council of Scientific and Industrial Research (CSIR), New Delhi, Government of India for providing the research fellowship. We acknowledge the Distributed Information Sub-Center (DISC) at the Institute of Life Sciences, Bhubaneswar, for the computational facility. We are thankful to Professor B. Schink (Universitaet Konstanz, Germany) for etymological advice.

## Authors contribution

S.K.D. developed the concept, R.R.A.K. conducted the experiment. R.R.A.K. and S.K.D. analysed the data and wrote the manuscript. All of us read and approved the final manuscript. The authors declare no conflicts of interest.

## Supplementary file

**Table S1.** Cellular fatty acid composition of strain RR6 and the type strains of related *Halopseudomonas* species. Strains: 1. *Halopseudomonas* sp. RR6; 2. *Halopseudomonas pachastrellae* CCUG 46540^T^ (Data from Wei et al. 2018); 3. *Halopseudomonas gallaeciensis* V113^T^ (Data from Mulet et al. 2018). ND, not detected.

| Fatty acid | 1 | 2 | 3 |
| --- | --- | --- | --- |
| 10:00 | 1.3 | 0.3 | ND |
| 10:0 3OH | 4.55 | 2.9 | 5.0 |
| 12:0 anteiso | 1.46 | ND | ND |
| 12:00 | 8.65 | 7.8 | 9.3 |
| 13:0 anteiso | 3.74 | ND | ND |
| 12:0 3OH | 3.33 | 3.7 | 4.2 |
| 14:00 | 1.14 | 1.1 | 1.1 |
| 15:0 anteiso | 3.15 | ND | ND |
| 16:1 w7c/16:1 w6c | 15.86 | 24.1 | 29.3 |
| 16:00 | 18.26 | 19.2 | 14.8 |
| 17:0 anteiso | 2.28 | ND | ND |
| 17:1 w8c | 0.22 | 1.4 | ND |
| 17:1 w6c | 0.33 | 1.1 | ND |
| 17:00 | 0.34 | 0.9 | ND |
| 18:1 w7c and/or 18:1 w6c | 28.32 | 34.4 | 33.4 |
| 18:00 | 1.14 | 0.9 | 1 |

**Figure S1.**
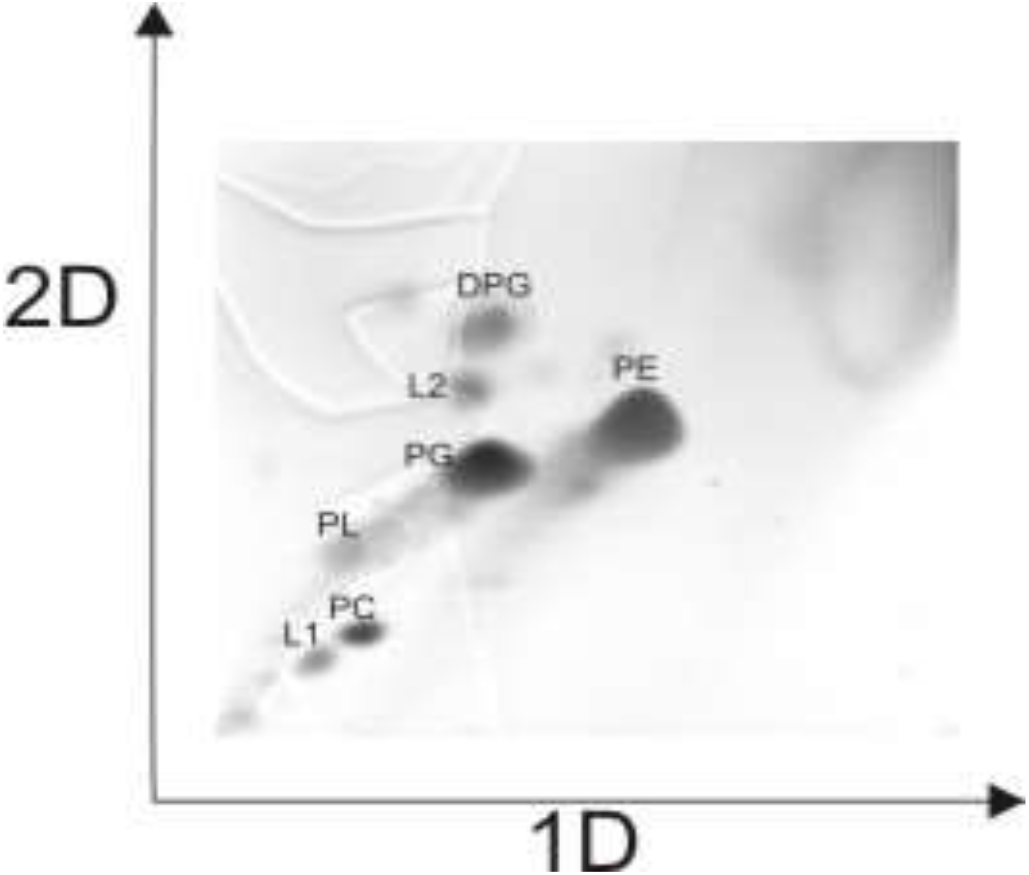
Two-dimensional thin-layer chromatographic profile of polar lipids of *Halopseudomonas s*train RR6. Abbreviations: Phosphatidylglycerol (PG), Diphosphatidylglycerol (DPG), Phosphatidylethanolamine (PE), Phosphatidylcholine (PC), unidentified phospholipid (PL) and unidentified lipids (L1-L2).

**Figure S2.**
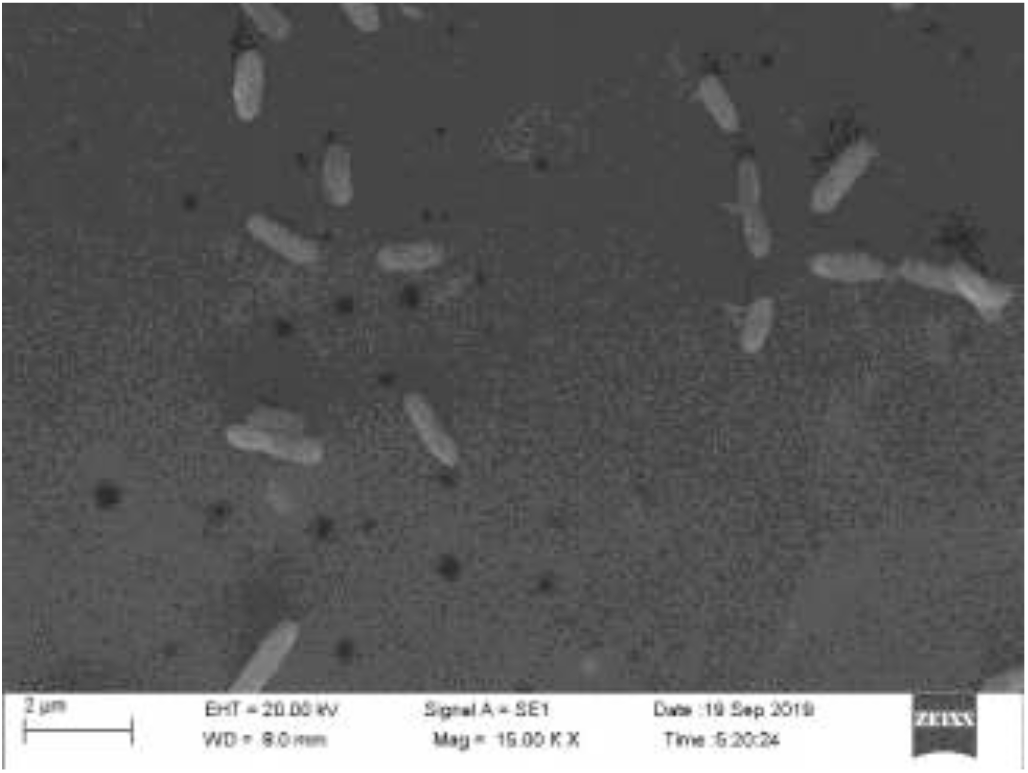
Scanning electron microscopic image of strain RR6

